# Supporting early detection of biological invasions through short-term spatial forecasts of detectability

**DOI:** 10.1101/2024.06.12.598508

**Authors:** César Capinha, António T. Monteiro, Ana Ceia-Hasse

## Abstract

Early detection of invasive species is crucial to prevent biological invasions. To increase the success of detection efforts, it is often essential to know the phenological stages in which the invasive species are found. This includes knowing, for example, if invasive insect species are in their adult phase, invasive plants are flowering, or invasive mammals have finished their hibernation. Unfortunately, this kind of information is often unavailable or is provided at very coarse temporal and spatial resolutions. On the other hand, opportunistic records of the location and timing of observations of these stages are increasingly available from biodiversity data repositories. Here, we demonstrate how to apply these data for predicting the timing of phenological stages of invasive species. The predictions are made across Europe, at a daily temporal resolution, including in near real time and for multiple days ahead. We apply this to detectability-relevant phenological stages of four well-known invasive species: the freshwater jellyfish, the geranium bronze butterfly, the floating primrose-willow, and the garden lupine. Our approach uses machine learning and statistical-based algorithms to identify the set of temporal environmental conditions (e.g., temperature values and trends, precipitation, snow depth, and wind speed) associated with the observation of each phenological stage, while accounting for spatial and temporal biases in recording effort. Correlation between predictions from models and the actual timing of observations often exceeded values of 0.9. However, some inter-taxa variation occurred, with models trained on several thousands of observation records performing consistently better than those based on a few hundred records. The analysis of daily predictions also allowed mapping EU-wide regions with similar phenological dynamics (i.e., ‘phenoregions’). Our results underscore the significant potential of opportunistic biodiversity observation data in developing models capable of predicting and forecasting species phenological stages across broad spatial extents. This information has the potential to significantly improve decision-making in invasion surveillance and monitoring activities.

## Introduction

Invasive alien species are a major environmental problem, severely impacting biodiversity, economies and public health (IPBES 2023). As human activities continue to transport and introduce alien organisms outside native regions (Hulme 2021; Capinha et al., 2023), the number of new invasions is expected to grow (Seebens et al. 2021), increasing the diversity and magnitude of their impacts. Because of this, successful invasion prevention efforts are key for safeguarding biodiversity, human livelihoods, and well-being (Vilà and Hulme 2017; IPBES 2023). In this regard, it is particularly important to detect non-native species in the early stages of the invasion process, as it significantly improves the effectiveness of control measures (Tobin et al. 2014; Larson et al. 2020; Martinez et al. 2020).

Current efforts in the surveillance and early detection of alien species encompass a large diversity of approaches, including camera and chemical traps, eDNA analysis, remote sensing, and visual surveys conducted by experts and citizen scientists (Larson et al. 2020). Each of these approaches has distinct strengths and limitations, and optimal outcomes are likely achieved through the integration and assimilation of their collective data (Larson et al. 2020; Fricke et al. 2023). A critical factor in enhancing the success and cost-effectiveness of most of these approaches is identifying the most suitable timing of the year for implementation. For instance, remote sensing often targets periods of higher species conspicuity, such as plant flowering or increased greenness, to enhance detection accuracy (e.g., Wijesingha et al. 2020). Similarly, trap-based surveillance programs are often tailored to specific life stages of the target species, and the deployment of these traps aims to coincide with the expected timing of these stages (Nguyen et al. 2024). Likewise, visual-based field surveys – either performed by experts or citizen scientists – also greatly benefit from knowledge about current or near future levels of species detectability (e.g., the timing of occurrence of the adult phase of an insect species, or of flower blooming), sustaining the development of surveys and monitoring activities when detectability is at its peak (Pocock et al. 2023).

Despite the importance of understanding the optimal timing for surveillance and early detection, information on species detectability levels is often unavailable, inadequate, or of limited value. For most invasive species, including highly problematic ones, the available information on these levels typically consists of dates of relevant life cycle stages observed in other regions (e.g., EFSA et al. 2020), or the months or seasons when these stages are typically observed (Flajšman et al. 2019). However, this type of information can overlook significant inter-regional and inter-annual differences resulting from the natural variation of drivers of phenology, such as temperature and precipitation (Godoy et al. 2009). Some exceptions exist for species for which phenological models have been developed. These models, whether process-based or data-driven, have yielded successful predictions of phenology (e.g., Barker et al. 2020; Reznik et al. 2022), thereby supporting invasion surveillance efforts (e.g., Taylor et al. 2020). However, they also suffer from notable limitations in terms of application and development for high numbers of invaders, i.e., ‘taxonomic scalability’. This is because process-based models largely depend on in-depth knowledge of the species’ eco-physiology, which can be costly and resource-intensive to acquire and is often only available *a priori* for a limited number of extensively studied species (Chuine and Régnière 2017). On the other hand, most data-driven phenological modelling approaches rely on long-term observation data, such as phenological time series (e.g., Taylor and White 2020), and these data are typically limited in terms of spatial and taxonomic coverage (Park et al. 2021). These limitations significantly hinder the application of such approaches to a broad range of taxa.

Recently, we have demonstrated how temporally and spatially discrete biodiversity observation data, widely available from popular online repositories such as the Global Biodiversity Information Facility (GBIF: https://www.gbif.org/) or iNaturalist (https://www.inaturalist.org/), can be used to estimate the timing of ecological phenomena across regions (Capinha et al. 2024). This approach is based on the concept of the phenological niche (Post 2019) and, in simple terms, involves using these data to represent the set of temporal environmental conditions under which an ecological phenomenon of interest (such as a species’ phenological stage) occurs. From a practical standpoint, this can be achieved by applying statistical or machine learning models to identify the ‘envelope’ of temporal environmental conditions associated with the observation of the phenomenon of interest. Once calibrated to perform this identification, the models can then be coupled with environmental predictor data (such as spatial time series of meteorological variables) and used to predict the probability of the phenomenon occurring over time (e.g., each day) and across regions.

Our previous work (Capinha et al. 2024) focused on demonstrating the conceptual and applied feasibility of this approach. Here, we aim to specifically highlight its potential for supporting efforts of early detection and monitoring of invasive species. This is done by demonstrating its use for predicting the timing of phenological stages of relevance for field surveying, at a daily resolution and for several days in advance, across Europe. We examine four well-known invasive species in Europe, offering different levels of observation data availability, and explore model accuracy, uncertainty, and the regionalization of predicted phenological patterns. Ultimately, we demonstrate that this approach has significant potential to inform invasion surveillance efforts while also holding strong taxonomic scalability.

## Methods

### Collation of observation record data

We focus on four alien species that are established in Europe: the freshwater jellyfish (*Craspedacusta sowerbii*), the geranium bronze butterfly (*Cacyreus marshalli*), the floating primrose-willow (*Ludwigia peploides*) and the garden lupin (*Lupinus polyphyllus*). The levels of visual detectability for these species change considerably throughout the year. The freshwater jellyfish is presumed to be native to regions of Asia and has been introduced in most continents of the world. However, its alien distribution remains poorly known, largely because the most visible part of its life cycle involves small medusae that appear for only a few months each year (Marchessaux et al. 2021). The geranium bronze is a small butterfly native to Southern Africa and currently invading parts of central and southern Europe. Like most insects, its adult (butterfly) stage has higher visibility due to increased mobility, and conspicuous colours. The floating primrose-willow is an aquatic plant native to Oceania and the Americas, with invasive populations in countries of central and southern Europe. This species produces bright yellow flowers, which facilitate its identification among surrounding vegetation (Booy et al. 2015). Finally, the garden lupin is a plant native to western North America that is now widespread in many temperate regions of the world, including Central and Northern Europe. This species produces prominent flower spikes (often violet, but sometimes also pink or white) that greatly facilitate its detection, including for remote sensing (Wijesingha et al. 2020).

Following our previously described framework (Capinha et al. 2024), we collected observation records supported by photographs for each of these species from GBIF. The records had to be made between 2016 and 2022 (a 7-year period matching the temporal extent of the environmental predictors used – see below) and provide the full date of observation (i.e., day, month, and year). The GBIF is a major online repository of biodiversity observation data, including from well-known and highly participated citizen-science projects (e.g., iNaturalist.org and observation.org), which typically provide supporting visual media. Because the number of records obtained from GBIF for the freshwater jellyfish was low, we also searched for observation records of this species in additional sources, particularly the USGS Nonindigenous Aquatic Species portal (https://nas.er.usgs.gov).

We visually checked the photographs supporting each observation record of the four species. Following this, we kept only the records referring to medusae of the freshwater jellyfish, the butterfly stage of geranium bronze, and the flowering stages of floating primrose-willow and the garden lupin. Records with images suggesting that the specimens were under human-care (e.g., garden lupin in gardens) were excluded. Likewise, we also excluded GBIF records where the observation date was the first day of the month and the observation time was ’00:00:00’. These are typically records where only the month and year of observation are known, and the first day of the month is assigned by default, i.e., the full date of the record may not be precise (Belitz et al. 2023). In total, we obtained 240 records for freshwater jellyfish medusae, 3,879 for geranium bronze butterflies, and 1,688 and 10,345 records for floating primrose-willow and garden lupin flowers, respectively.

### Environmental drivers

We collected time series of global-scale maps representing daily conditions of maximum, minimum and mean temperature, accumulated precipitation, wind speed and accumulated snow. These factors are expected to be drivers of the timing of occurrence of the species life stages of interest, according to previous research (Favilli and Manganelli 2006; Ludewig et al. 2022; Marchessaux et al. 2022). The data were collected from the Global Forecast System (GFS), a large-scale NOAA weather forecast model (https://www.ncei.noaa.gov), covering the period between 15^th^ January 2015 (the first day the data is available) to 31^st^ December 2022. GFS data is provided at 0.25° spatial resolution, for multiple hourly intervals and for each model run that takes place at 00, 06, 12, and 18 UTC daily. The daily-resolution maps were obtained by aggregating the first six-hour values provided by each model run. The temporal immediacy of this period in relation to the timing of model runs results in highly accurate weather forecasts (NOAA, 2022). Models employing these data achieve results comparable to those using climate reanalysis data - traditionally for ecological forecasting - like ERA5 (Capinha et al. 2024). Crucially, NOAA also offers real-time access to GFS data, including weather forecasts extending several days into the future, meeting our objective to deliver predictions in real-time and for short-term forecasting.

### Spatial bias removal

We implemented a set of procedures to minimise potential spatial and temporal biases in the observation data. Spatial bias refers to unequal numbers of records in distinct regions, which can lead to model responses being ‘dominated’ by the patterns occurring in oversampled regions. Temporal observation bias arises from unequal levels of recording effort over time, confounding the actual temporal signal of phenological events.

To address these biases, we followed the procedures we proposed earlier (Capinha et al. 2024). Specifically, and to minimise spatial bias, we kept only one record located in the same grid cell and having the same date of observation. Next, we also downsampled observations in oversampled regions. For this purpose, we used a reference grid of 250 ⨉ 250 km squares, for which we identified squares that were upper outliers in terms of record count (i.e., *n* > third quartile + 1.5 * interquartile range). For these areas, we randomly subsampled a number of records equal to the outlierness threshold.

### Temporal bias removal

Our framework includes an optional procedure to minimise temporal bias, named ‘benchmark taxa approach’ (Capinha et al. 2024). The rationale is to use taxa that remain similar in their visual appearance along the year as an indicator of variation in levels of activity of recorders (e.g., citizen scientists). The temporal variation in the frequency of records for these taxa is related to variables expected to mediate levels of recording effort (e.g., days of the week, months of the year and weather conditions) by means of a statistical model such as a generalised linear model. Based on the relationships identified, the temporal biases in records of the phenomenon of interest can be minimised by a subsampling procedure, where records made in periods of higher levels of recording intensity receive a lower probability of being selected for model development. We demonstrated this approach previously, and its application delivered similar performance to the models without using it. However, it is not clear if this outcome can be expected in the generality of phenomena. Hence, here we performed all the analyses using event observation data with this correction (described in Text S1 of Supplementary Materials), and without it. The results were similar for both cases (see results), hence those of the former are provided in the supplementary materials.

### Environmental characterization of records

We next characterized the meteorological conditions preceding each event record. We used a total of 67 features representing multiple features of temperature (e.g., maximum, minimum and mean values, growing degree days and cold accumulation), accumulated precipitation, accumulated snow, and mean wind speed for distinct preceding periods (see full list in Table S1).

Additionally, we assembled a second set of records aimed at representing the meteorological conditions that are generally available in the location of each of the events (i.e., the background environmental conditions). This was performed using ‘temporal pseudo-absences’ (Capinha et al. 2024), which correspond to records having the same geographical coordinates as event records, but with dates drawn at random within the temporal range of the event data. A total of 12 pseudo-absence records were generated from each event record and each was characterised using the same set of 67 environmental features.

### Model training and evaluation

Prior to model fitting, we tested for the presence of multicollinearity among the predictors. For this purpose, we measured their variance inflation factor (‘VIF’) and excluded any predictor with a VIF value above 10 (Gareth et al. 2021). We then trained three machine learning algorithms: random forests (RF), boosted regression trees (BRT) and generalised linear models with lasso regularization (GLM-lasso), in distinguishing the conditions associated to the phenological stages of interest and those represented by the temporal pseudo-absences. These algorithms were selected because they are commonly used for ecological modelling and prediction, and often rank amongst the best performing, including when transferred to new spatial settings (Zhang et al. 2019; Valavi et al. 2023).

The implementation of these models was performed in R (R Core Team, 2024), using the ‘randomForest’ package for RF (Liaw and Wiener 2022), the ‘dismò package for BRT (Hijmans et al. 2017), and ‘glmnet’ for GLM-lasso (Hastie et al. 2021). We optimized several parameters within these functions to improve model fitting. Random forests used 2,000 individual trees (instead of the default 500) to increase the chances of the relatively large number of predictor variables and samples being adequately represented in the final ensemble. For the BRT models, the number of trees in each ensemble was automatically determined by the ‘gbm.step’ function of the ‘dismò package, with a tree complexity of 3 (allowing for interactions among predictor variables) and a learning rate of 0.005. Additionally, for all three algorithms, we addressed the class imbalance in the data (i.e., one event observation record for every 12 pseudo-absence records), which could skew models toward overpredicting the dominant class (pseudo-absences). For GLM-lasso and BRTs, this was made through a weighting parameter (‘weights’ for GLM-lasso and ‘site.weights’ for BRTs), assigning class-proportional weights to each sample. For Random Forests, the adjustment was made using the ‘sampsizè parameter of the ‘randomForest’ function, ensuring that each individual tree used an equal number of event observation records and temporal pseudo-absences.

To evaluate the predictive performance of the models, we used a leave-one-year-out cross-validation procedure. This involved excluding the data from one year for model calibration and using it to assess the predictive ability of models trained on the remaining years. The procedure was iterated so that the data from each year served as an evaluation set. To measure the models’ performance, we used the Boyce index, initially proposed for species distribution models (Hirzel et al. 2006). The Boyce index measures the correlation between the frequency of observation records and predicted probability values. In the context of our work, a strong positive correlation implies that event observations are concentrated in periods predicted with higher probability values, whereas correlations near zero indicate predictions akin to those expected from a model that assigns probability values randomly throughout the year. We preferred this metric over discrimination-based metrics, like the commonly used area under the receiver operating characteristic curve (Pontius and Parmentier 2014), because these latter metrics would involve assessing the misclassification of temporal pseudo-absences. However, in the context of our work these pseudo-absences capture the environmental variation that is available, including periods that are potentially suitable for the occurrence of our focal phenological stages. This becomes particularly critical for models of phenological stages with extended seasons of occurrence, where temporal pseudo-absences during suitable periods are more likely. Consequently, this would likely result in an underestimation of the models’ true predictive performance due to the misclassification of pseudo-absences.

We performed the Boyce index calculations using the *ecospat.boyce* function from the *ecospat* R package (Broennimann et al. 2015), employing the Pearson correlation coefficient as a metric. The measurements were performed for the three algorithms and for an ensemble of these predictions corresponding to their average.

### Real-time, days-ahead forecasts of event probability

The ability to predict species’ phenological stages several days in advance can guide decision on the optimal the timing of invasion detection actions (Barker et al. 2020; Crimmins et al. 2020). Consequently, an important objective of our work was to use the models developed to provide real-time phenological forecasts for several days into the future. We performed this based on GFS weather forecasts and for up to 9 days into the future, a forecast horizon that is broad enough to guide short-term surveying decisions, but which is not too close to the limit of the GFS forecast horizon (16 days), allowing to accommodate technical challenges such as server downtime and decreased temporal resolution of GFS data after 10 days. However, an important question is whether the phenology forecasts lose accuracy as they extend further into the future, and if so, to what extent. To address this, we calculated the Boyce index for phenology forecasts derived from GFS weather data produced immediately before the target day (i.e., the 18:00 UTC run of the day before). We then compared these with forecasts based on weather data generated 3, 6, and 9 days in advance. This assessment covered a 10-month period from June 1, 2023, to March 31, 2024, corresponding to the timeline from the real-time deployment of the forecasting models to the writing of this work. As evaluation data, we gathered observation records for this period from the GBIF, keeping only those that represented the life stages of interest, and performing the same initial data cleaning procedures as for calibration data (i.e., removing records without full date attributes and duplicates in space and time). Only records in Europe were considered, matching the geographical focus of the work (i.e., where the four species are invasive). Observation records for *L. polyphyllus* for this period were highly voluminous (>19,000 records). To reduce the time resources needed to visually identify the life stage of each observation, a subset of 1000 randomly selected records were considered for processing.

### Mapping phenoregions

Identifying regions with similar year-to-year phenological patterns (“phenoregions”; White et al. 2005) allows defining areas where invasion surveillance efforts could be implemented in the same periods. To identify these regions, we used the phenology predictions from RF models at 5-day intervals for the 7 years period (from 2016 to 2022), across Europe. We used the predictions from this algorithm because it consistently performed well, ranking as the best or second-best across all species (see Results). To minimize the risk of extrapolation errors, i.e., making predictions for conditions far from those represented in the training data, we only made predictions for regions having Koppen climate classes (collected from Beck et al. 2018), where each of the phenological stages had at least 5 observation records. Next, we applied a k-means algorithm to cluster regions based on the temporal variation in predicted values, using the ‘elbow’ method to determine the optimal number of clusters (Syakur et al., 2018). For each identified region, we then calculated the mean and standard deviation of inter-annual variation.

## Results

### Predictive performances

Overall, the predictions from models (Fig. 1) achieved high performances. The predictions of imago-stage of the Geranium bronze attained a cross-model median correlation with the timing of observations of 0.95. GLM-Lasso achieved the highest median performance (*r* = 0.97), followed by RF (*r* = 0.96), multi-model ensemble (*r* = 0.95) and BRT (*r* = 0.94) (Fig. 2). Pairwise Kruskal-Wallis tests indicate non-significant differences in these performances (i.e., *p* ≥ 0.05; Table S2). Likewise, the flowering timings of the floating primrose-willow and the garden lupin were also well captured by the models, with median cross-model correlations of 0.94 and 0.91, respectively. For the floating primrose-willow, the multi-model ensembles, RF, and GLM-Lasso models had the best performance (median *r* = 0.94), with BRT performing slightly worst (median *r* = 0.88). However, as before these differences were not statistically significant (Kruskal-Wallis *p* ≥ 0.05; Table S2). In contrast, for the garden lupin, RF displayed a statistically significant superiority (median *r* = 0.96; Kruskal-Wallis *p* ≤ 0.05; Table S2), over remaining algorithms. BRT was second best (median *r* = 0.92), followed by multi-model ensemble and GLM-Lasso (median *r* for both = 0.88), but with no statistically significant differences among these (Kruskal-Wallis *p* ≥ 0.05; Table S2). Predictive performance for the medusae stage of the freshwater jellyfish was the lowest but still reasonable – with a median correlation of 0.73 across models (Fig. 2). Predictions from the ensemble approach demonstrated a substantially higher performance (median *r* = 0.82), followed by RF (median *r* = 0.79), GLM-Lasso (*r* = 0.64), and BRT (*r* = 0.42), but these differences were also not statistically significant (Table S2).

**Figure 1.**
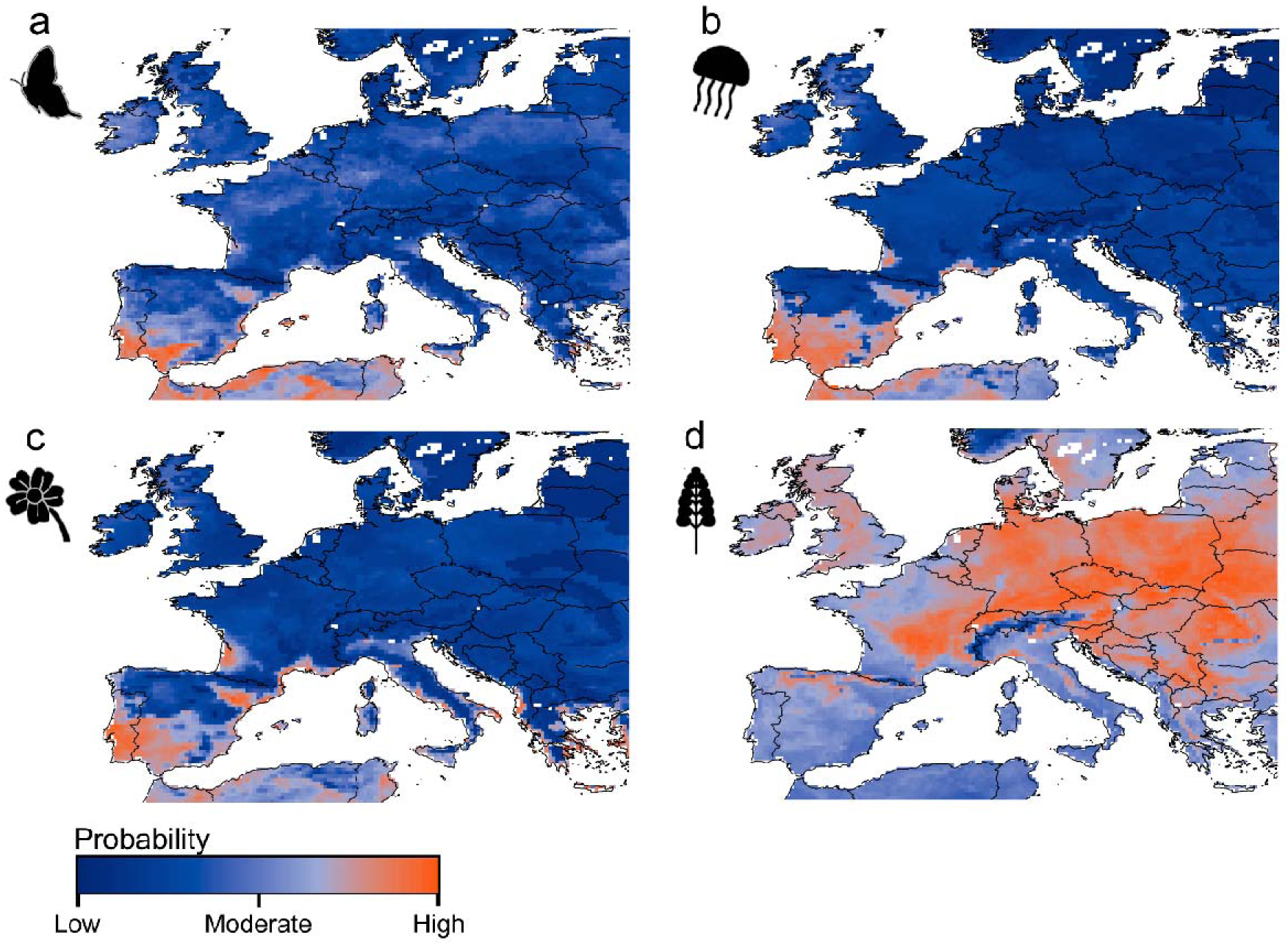
Examples of daily predictions obtained from the modelling approach. These represent the probability of occurrence for each of the four modelled phenological stages (imago-stage of the Geranium bronze [a]; medusae of the freshwater jellyfish [b], flowering of the floating primrose-willow [c] and flowering of garden lupin [d]), for July 1st, 2023. Predictions were obtained using the random forest algorithm trained with observational data corrected for spatial bias.

**Figure 2.**
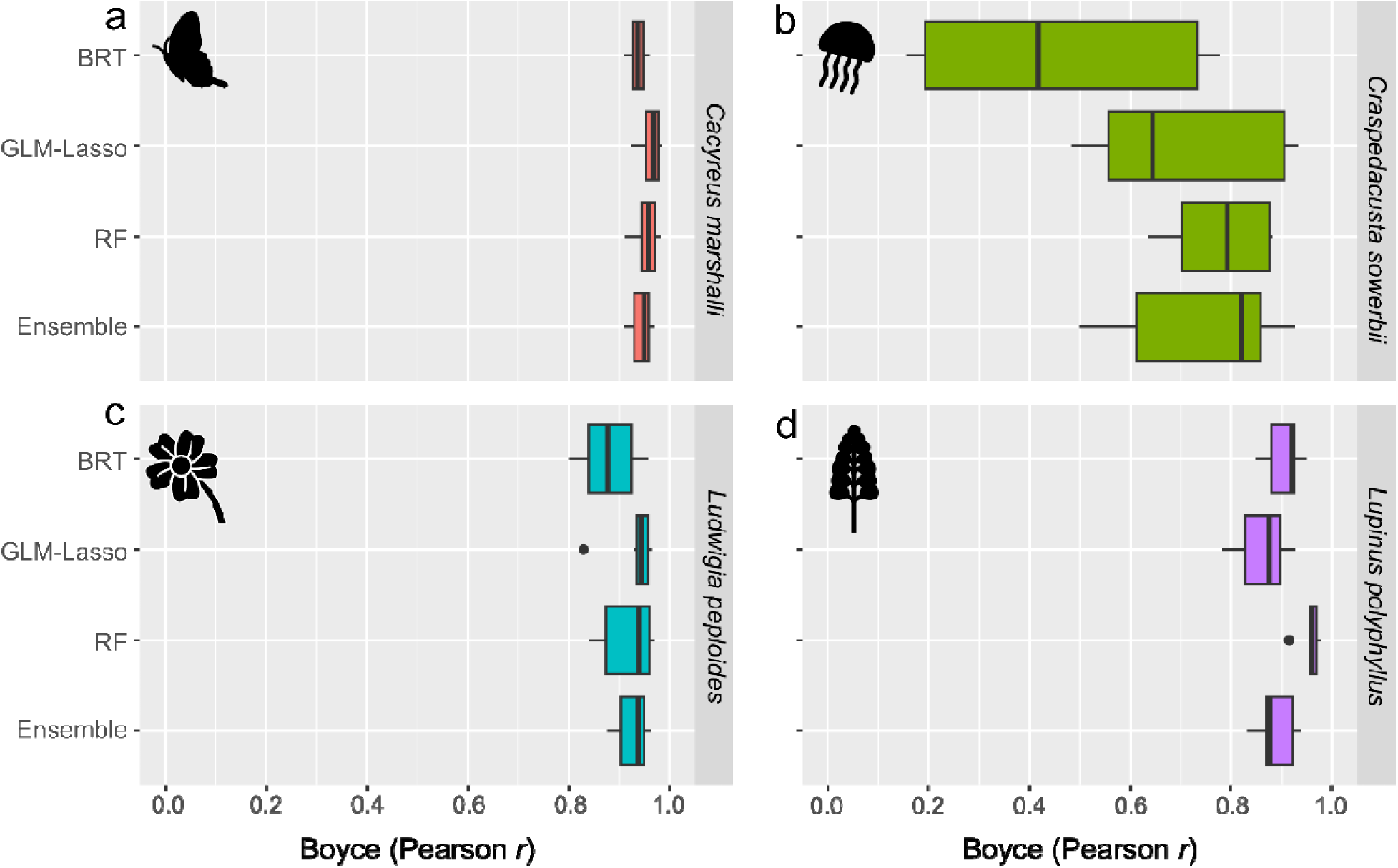
Results of Boyce index corresponding to Pearson correlation values between predicted probabilities of event occurrence and the frequency of event observation records of the imago stage of the Geranium bronze butterfly (a), medusae of the freshwater jellyfish (b), and the flowering of floating primrose-willow (c) and the garden lupin (d). The boxplots represent the variation of correlation values assessed for 7 years (2016 to 2022), using three modelling algorithms (boosted regression trees, BRT; generalised linear models with lasso regularization, GLM-Lasso; random forest, RF) and an ensemble of previous algorithms (Ensemble), trained with observation data corrected for spatial bias.

Relevantly, models trained with data addressing both spatial and temporal biases showed similar predictive performances as those addressing only spatial biases (Fig. S1). Specifically, the predictions of imago-stage of the Geranium bronze retained the highest cross-model median correlation with the timing of observations (0.95), followed by the garden lupin (0.91), the floating primrose-willow (0.90) and the freshwater jellyfish (0.67). Performances for the Geranium bronze were highest for GLM-Lasso, RF and multi-model ensemble (median *r* = 0.95), followed by BRT (median *r* = 0.90). For the floating primrose-willow, the best predictions were achieved by multi-model ensembles and RF (median *r* = 0.91), followed by GLM-lasso (median *r* = 0.90) and BRT (median *r* = 0.87). For the garden lupin, the best performances were achieved by RF (median *r* = 0.97), followed by multi-model ensembles and BRT (median *r* = 0.90) and by GLM-Lasso (median *r* = 0.88). Finally, for the freshwater jellyfish the best performances were achieved by RF and GLM-Lasso (median *r* = 0.68), followed by multi-model ensembles (median *r* = 0.66) and BRT (median *r* = 0.44). As for models using observation data corrected for spatial bias only, the only statistically significant difference among algorithms was verified for the garden lupin, with RF significantly outperforming remaining ones (Table S2).

We also assessed the performance of days-ahead forecasts across Europe over a 10-month period (Fig. 3; Fig. S2). For the freshwater jellyfish and the floating primrose-willow, the observation records used for model evaluation during this period were relatively limited (25 and 42, respectively), while they were more abundant for the Geranium bronze (1,187) and the garden lupin (126 records from a random sample of 1,000 processed records). Overall, the forecasts demonstrated a high correlation with the timing of observed life stages modelled, with values generally close to 0.8 or higher (Fig. 3; Fig. S2). Additionally, there was no apparent pattern of decreased model performance as the forecast horizon extended (e.g., from 1 day ahead to 9 days ahead).

**Figure 3.**
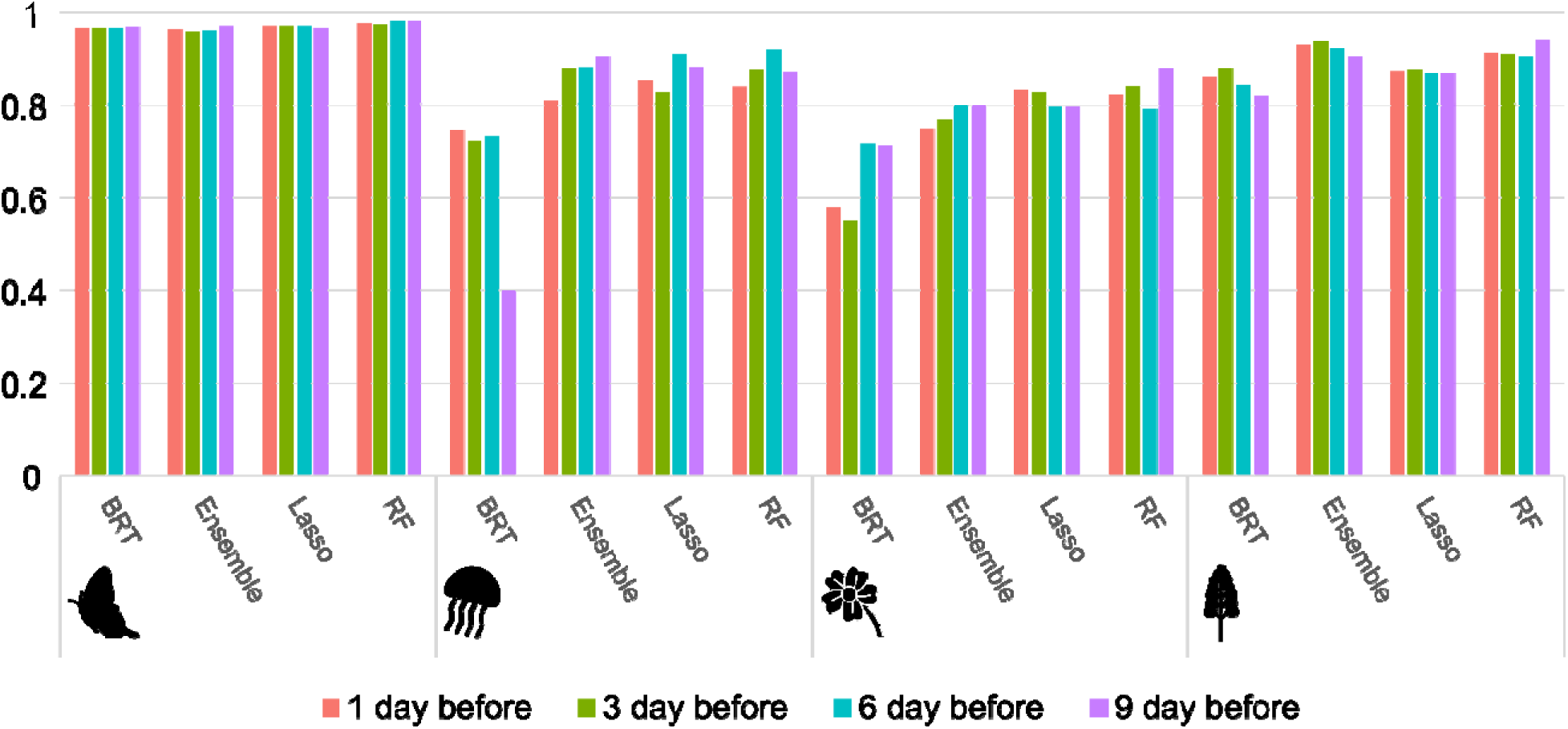
Boyce Index values for forecasts made 1, 3, 6, and 9 days in advance. These values correspond to the Pearson correlation coefficient between predicted probabilities of event occurrence from July 2023 to March 2024 and the frequency of event observations recorded during the same period. The values are reported for three modelling algorithms—boosted regression trees (BRT), generalized linear models with lasso regularization (Lasso), and random forest (RF)—as well as an ensemble of these algorithms (Ensemble), all trained with observation data corrected for spatial bias.

### Regions of similar phenology (‘phenoregions’) and associated temporal patterns

Using the predictions of the random forest algorithm trained with observation data corrected for spatial bias, we identified five phenoregions for the Geranium bronze butterfly in Europe (Fig. 4a), with the peak occurrence ranging from early June to late November in the southernmost region (Fig. 4b). The remaining regions have peak occurrences between August and September, with northern latitudes having a shorter season of occurrence, and the opposite being true for southern latitudes. Recorded observations of butterflies of this species are concentrated in the summer (Fig. 4c), especially in the mid- to higher latitude regions, in agreement with the predictions.

**Figure 4.**
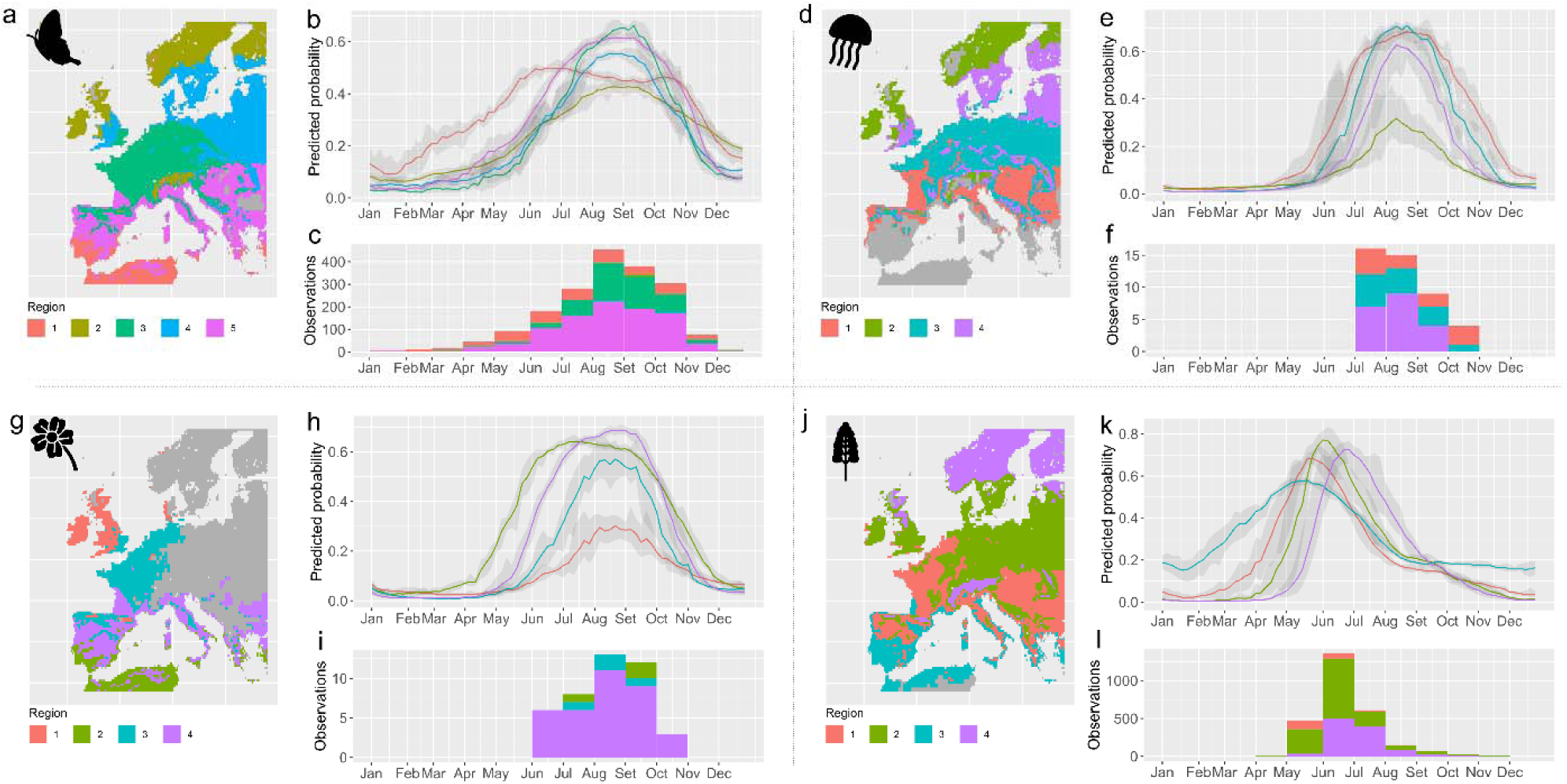
Regional patterns of predicted and observed timings of the emergence of the imago stage in the Geranium bronze butterfly (panels a-c), the occurrence of medusae in the freshwater jellyfish (panels d-f), and the flowering phases of floating primrose-willow (panels g-i) and garden lupin (panels j-l). The maps display areas having similar phenological dynamics (‘phenoregions’), based on daily projections at 5 days intervals from 2016 to 2017. Areas in grey represent environmental conditions far from those represented in the training data and were not considered in the analysis to minimise risks of model extrapolation. Time series depict the inter-annual mean probabilities of occurrence of each event, along with their standard deviations (grey shading), for each region throughout the year. Histograms show the monthly frequency of effectively observed occurrences of each life stage within each phenoregion. Predictions and phenoregion delineations were made also for areas where the species have not yet been recorded, leading to observation records being absent from the histograms for certain regions.

Predictions of the timing of occurrence of medusae of the freshwater jellyfish were clustered into four phenoregions, peaking between late August and September. However, southern regions exhibit higher probabilities of occurrence over substantially longer periods, and conversely, shorter periods are predicted for northern regions. Medusae observations take place in the months of predicted peaks, except for the mid-latitude region covering most of Central Europe, where they concentrate in September and November— a period when the predicted probabilities are already declining.

For the floating primrose-willow four regions were identified (Fig. 4g), with the southern regions showing earlier and more extended flowering periods (Fig. 4h). The timing of observational data matches well with the predictions for the region including most of Mediterranean Europe, although very few records are available for the other regions (Fig. 4i).

Predicted timings of flowering for the garden lupin were classified into four phenoregions (Fig. 4j), with probabilities peaking from early May to late June and occurring earlier in southern regions (Fig. 4k). Observational data also largely agree with the predicted patterns, especially for the two northernmost regions, which have most records (Fig. 4l).

Importantly, identified phenoregions and associated temporal patterns were strikingly similar to those obtained based on models calibrated with temporally unbiased observation data (Fig. S3). This further suggests the robustness of predictions to temporally varying levels of recording effort.

## Discussion

Efforts for the early detection of biological invasions benefit greatly from knowledge about temporal variation in detectability levels of the species of concern. Here, we have demonstrated that temporally discrete biodiversity observation data, as made widely available in public repositories such as GBIF and extensively contributed by citizen science (Bonney et al. 2021), can be used to estimate the timing of species life stages of relevance for detectability, including across broad spatial regions (e.g., continents) and at very high temporal resolution (e.g., daily).

Our results demonstrate a varying ability of trained models to identify the temporal environmental variation associated with observing the life stages of interest. For three of the life stages modelled, the agreement between predictions and the timing of observation was, across models, very high, with median correlations regularly above 0.9. However, for one of them (the medusae stage of the freshwater jellyfish), the performance was lower (though still high; cross-model correlations = 0.73 and 0.67). This lower performance coincides with the life stage represented by the lowest number of observation records available (few hundreds), significantly fewer than those available for the remaining life stages (in the order of thousands). This strongly suggests that, as with most modelled phenomena, the size of the training data can be a limiting feature. Indeed, the predictions allowed by our approach, which can be made daily and over wide geographical areas, involve a high dimensionality of conditions in the prediction space resulting from the multiple states of preceding conditions for each environmental variable and their joint combination. Hence, it is a reasonable expectation that the calibration data should be necessarily large in number to represent this variability; otherwise, extrapolation will occur, and the uncertainty of predictions will be higher and possibly also less accurate. Nonetheless, it is still very positive to verify that sample sizes on the order of magnitude of a few thousands (as available for the remaining species), consistently deliver very good predictions. Indeed, the number of invasive species for which life stages of relevance for early detection and monitoring are represented by similar amounts of records can be presumed to easily reach several thousands, considering the enormous volumes of biodiversity data becoming available each year, particularly through citizen science. This highlights the potential taxonomic scalability of our approach.

It is well-known that multi-sourced opportunistic biodiversity observation data, as used to train our models, can suffer from substantial spatial and temporal bias, often hindering efforts of use for prediction (Isaac and Pocock, 2015). We applied a set of procedures to minimise the effect of these biases. Specifically, we aimed at correcting for the temporal bias component (see also Capinha et al. 2024). However, we also found that models having this correction and those without it are largely equivalent in terms of predictions. This was also verified in our previous work demonstrating the general modelling workflow (Capinha et al. 2024). As discussed there, we interpret these results as virtue of the way models are trained. Conceptually, our models contrast two point clouds in the environmental space, one representing the conditions associated to the occurrence of the life stage of interest and the other the generality of conditions available in the regions of its occurrence. Because the number of random records used to represent available conditions is generated at a fixed proportion of the observation records, observation records from periods of lower recording effort will have an equivalent relative influence in model training to those from periods of higher effort, (which will have, in turn, a higher number of contrasting random records). This does not mean that predictions made for the least represented conditions will be as accurate as those made for more populated spaces of the environmental space. Rather, it means that having more populated spaces of the environmental space will not direct the models’ inference towards these conditions. Ultimately, these low-sampled environmental spaces raise issues of extrapolation, and operational predictions from the models should be avoided here. For instance, in the phenoregion maps, we minimised extrapolation issues by only making predictions for Köppen climate regions having five or more observation records of the life stage modelled. This number was selected subjectively, and its robustness in excluding areas of extrapolation should be considered more thoroughly in future work.

Our approach is also capable of forecasting the probability of occurrence of the life stages assessed, in real time and for several days into the future. This capacity is perhaps the most impactful aspect of our work for practical applications. Providing these forecasts daily and across extensive areas (such as the European continent in this case) could support a variety of decision-making processes related to the timing and efficacy of invasive species detection efforts. Relevantly, we also observed that the performance of forecasts remains largely stable over the forecast horizon considered. This consistency may result from the relatively brief time span considered (nine days) and the inherent temporal correlation among phenological events, which tend to unfold gradually and slowly over time (considering the daily temporal resolution used).

The identification of phenoregions (i.e., regions sharing similar phenological dynamics), as allowed through the spatial clustering of predictions from our framework, can be of great interest to support invasion surveillance and decision-making. Environmental managers are often left wondering which time of the year specific life stages of species will occur. Most of the information available (when available) is found in technical and scientific literature and typically indicates the months or seasons of occurrence at broad geographical resolutions, e.g., a country, group of countries, or a continent. For instance, a highly comprehensive recent work on invasive species in the forests of Europe (Veenvliet et al. 2017) highlights the months of expected occurrence of life stages with the highest visibility for each species across Europe as a whole. This information is much welcomed by managers and is often the best available for most species, but it well illustrates the difficulties of obtaining high temporal and spatial resolution data on the timing of phenological stages. The maps obtained with our approach go a step further, identifying these regions at the level of individual grid cells, with temporal resolutions as fine as daily, and by accounting for inter-annual variability.

In conclusion, our work demonstrates the potential of widely available, temporally discrete biodiversity observation data for estimating the timing of life stages relevant to invasive species detectability. With the increasing volume of media-supported biodiversity observation data being published, the number and diversity of invasive species for which these estimates can be produced are substantial. Furthermore, these estimates can be delivered at high spatial resolutions across wide areas, in real time and for several days into the future, providing timely decision support for numerous managers tasked with planning surveillance and early detection measures. Increasing the number of invasive species covered, while continuously refining these estimates, will likely contribute significantly to global efforts in the proactive prevention of biological invasions.

## Acknowledgements

The forecasts here described are implemented in real-time and can be accessed at https://www.natureforecast.org.

## Data Resources

Spatial weather data used for this work are publicly available online from National Centers for Environmental Prediction. Event observation data are available from USGS Nonindigenous Aquatic Species (https://nas.er.usgs.gov/queries/factsheet.aspx?SpeciesID=1068) and GBIF with DOIs: 10.15468/dl.3fve6q, 10.15468/dl.9drr85, 10.15468/dl.h5amhh, 10.15468/dl.krbguc, 10.15468/dl.2kbnxv, 10.15468/dl.ycjsyz, 10.15468/dl.7q6rke, 10.15468/dl.b9sze9 and 10.15468/dl.uh8apz. R code will be made publicly available on Zenodo (https://zenodo.org/) upon publication of the manuscript.

## Author Contributions

CC: Conceptualization, Methodology, Validation, Formal analysis, Resources, Data Curation, Writing - Original draft, Project administration, Funding Acquisition. AC and AM: Writing - Review and Editing.

