## Supplementary Information for "Supporting early detection of biological invasions through short-term spatial forecasts of detectability"

**Text 1 – Rationale and procedures used to address temporal recording bias.**

The rationale and procedures for addressing temporal recording bias are comprehensively described in Capinha et al. (2024). Here, we provide a concise overview, focusing on the methodological steps.

The approach involves using a ‘benchmark taxonomic group’, specifically Pinus spp. (i.e., pines), which are expected to exhibit temporal variability in record availability primarily due to observation effort rather than phenological changes. Observation records of Pinus spp. from 2016 to 2022 were used, undergoing the same data cleaning procedures as those for the life stages of interest (see main document). Additionally, a set of records with the same geographical coordinates but randomly generated dates within the years was created. These records represent a distribution of observations that would occur if they were made randomly over time. For both types of records (observations and randomly generated dates), calendar and weather predictors were extracted, including the day of the week, month, average temperature, total precipitation, and average wind speed. These predictors are expected to capture the most relevant factors influencing recording effort.

A generalized linear model (GLM) with a binomial error distribution was then used to relate the two classes of records (benchmark taxa observations, coded as 1, and randomly generated dates, coded as 0) to the calendar and weather predictors. The resulting model was applied to predict the level of sampling effort for the conditions associated with each record of the four life stage events being modelled. The predicted values represent the propensity for increased records due to more favourable conditions for observers.

Subsequently, a second data set of observation records was built for each event. Using data sets corrected for spatial bias, the probability of each original observation being included in the new data set was defined to be inversely proportional to the predicted level of observation effort (i.e., inverse probability sampling). This involved randomly selecting, with replacement, the same number of total records, where the probability of each record being selected was 1 minus the probability predicted by the model. Thus, records of species life stages made under less favourable conditions for observers had a higher chance of being selected, and vice versa. These datasets were used to calibrate models, correcting for both spatial and temporal bias in observation data.

| **Table S1.** List of the 67 features used to characterize temporal environmental conditions for observation and temporal pseudo-absence records. ‘T’ stands for the date of the observation records and the associated numbers represent preceding days. | | | | | | | | | | | | | |
| --- | --- | --- | --- | --- | --- | --- | --- | --- | --- | --- | --- | --- | --- |
| **Geography** |  |  | **Mean temperature** |  | **Minimum temperature** |  | **Maximum temperature** |  | **Precipitation** |  | **Snow depth** |  | **Wind speed** |
| Latitude |  |  | Mean T -1 to T -2 |  | Mean T -1 to T -7 |  | Mean T -1 to T -7 |  | Sum T -1 to T -2 |  | Mean T -1 to T -5 |  | Mean T -1 to T -3 |
| Longitude |  |  | Mean T -1 to T -5 |  | Mean T -8 to T -14 |  | Mean T -8 to T -14 |  | Sum T -1 to T -4 |  | Mean T -1 to T -15 |  | Mean T -1 to T -7 |
|  |  |  | Mean T -6 to T -10 |  | Mean T -15 to T -21 |  | Mean T -15 to T -21 |  | Sum T -5 to T -8 |  | Mean T -16 to T -30 |  | Mean T -8 to T -14 |
|  |  |  | Mean T -11 to T -15 |  |  |  |  |  | Sum T -9 to T -12 |  |  |  |  |
|  |  |  | Mean T -16 to T -20 |  |  |  |  |  | Sum T -13 to T -16 |  |  |  |  |
|  |  |  | Mean T -21 to T -30 |  |  |  |  |  | Sum T -1 to T -8 |  |  |  |  |
|  |  |  | Mean T -31 to T -40 |  |  |  |  |  | Sum T -9 to T -16 |  |  |  |  |
|  |  |  | Mean T -41 to T -50 |  |  |  |  |  | Sum T -17 to T -24 |  |  |  |  |
|  |  |  | Mean T -51 to T -60 |  |  |  |  |  | Sum T -25 to T -32 |  |  |  |  |
|  |  |  | Mean T0 to T -29 |  |  |  |  |  | Sum T -33 to T -47 |  |  |  |  |
|  |  |  | Mean T -30 to T -59 |  |  |  |  |  | Sum T -48 to T -62 |  |  |  |  |
|  |  |  | Mean T -60 to T -89 |  |  |  |  |  | Sum T0 to T -29 |  |  |  |  |
|  |  |  | Mean T -90 to T -119 |  |  |  |  |  | Sum T -30 to T -59 |  |  |  |  |
|  |  |  | Mean T -120 to T -149 |  |  |  |  |  | Sum T -60 to T -89 |  |  |  |  |
|  |  |  | Mean T -150 to T -179 |  |  |  |  |  | Sum T -90 to T -119 |  |  |  |  |
|  |  |  | Mean T -274 to T -364 |  |  |  |  |  | Sum T -120 to T -149 |  |  |  |  |
|  |  |  | Mean T -182 to T -273 |  |  |  |  |  | Sum T -150 to T -179 |  |  |  |  |
|  |  |  | Mean T -182 to T -364 |  |  |  |  |  | Sum T -180 to T -209 |  |  |  |  |
|  |  |  | Mean T0 to T -365 |  |  |  |  |  | Sum T -274 to T -364 |  |  |  |  |
|  |  |  | Growing degree days since 1st Julian day (baseline 0ºC) |  |  |  |  |  | Sum T -182 to T -273 |  |  |  |  |
|  |  |  | Growing degree days since 1st Julian day (baseline 7ºC) |  |  |  |  |  | Sum T -182 to T -364 |  |  |  |  |
|  |  |  | Growing degree days since 1st Julian day (baseline 15ºC) |  |  |  |  |  | Sum T0 to T -365 |  |  |  |  |
|  |  |  | Growing degree days of past 90 days (baseline 2ºC) |  |  |  |  |  |  |  |  |  |  |
|  |  |  | Growing degree days of past 60 days (baseline 2ºC) |  |  |  |  |  |  |  |  |  |  |
|  |  |  | Growing degree days of past 30 days (baseline 2ºC) |  |  |  |  |  |  |  |  |  |  |
|  |  |  | Growing degree days of past 30 to 15 days (baseline 2ºC) |  |  |  |  |  |  |  |  |  |  |
|  |  |  | Growing degree days of past 15 days (baseline 2ºC) |  |  |  |  |  |  |  |  |  |  |
|  |  |  | Growing degree days of past 7 days (baseline 2ºC) |  |  |  |  |  |  |  |  |  |  |
|  |  |  | Cold accumulation since 1st Julian day (baseline 5ºC) |  |  |  |  |  |  |  |  |  |  |
|  |  |  | Cold accumulation since 1st Julian day (baseline 10ºC) |  |  |  |  |  |  |  |  |  |  |
|  |  |  | Cold accumulation of past 30 days (baseline 5ºC) |  |  |  |  |  |  |  |  |  |  |

**Table S2.** Results from pairwise Kruskal-Wallis tests assessing significant differences in the performance of predictions from distinct modelling algorithms. Significant differences (in bold) are defined at α = 0.05). Three modelling algorithms (boosted regression trees, BRT; generalised linear models with lasso regularization, Lasso; random forest, RF) and an ensemble of previous algorithms (Ensemble) are compared. The results in the left panel refer to models calibrated using observation records corrected for spatial recording bias, while those on the right are corrected for both spatial and temporal biases.

| ***C. marshalii*** | BRT | Ensemble | Lasso |  | ***C. marshalii*** | BRT | Ensemble | Lasso |
| --- | --- | --- | --- | --- | --- | --- | --- | --- |
| Ensemble | 0.48 | - | - |  | Ensemble | 0.89 | - | - |
| Lasso | 0.23 | 0.33 | - |  | Lasso | 0.89 | 1 | - |
| RF | 0.33 | 0.48 | 0.48 |  | RF | 0.89 | 1 | 1 |
| ***C. sowerbii*** | BRT | Ensemble | Lasso |  | ***C. sowerbii*** | BRT | Ensemble | Lasso |
| Ensemble | 0.16 | - | - |  | Ensemble | 0.42 | - | - |
| Lasso | 0.26 | 1 | - |  | Lasso | 0.22 | 0.8 | - |
| RF | 0.16 | 0.8 | 1 |  | RF | 0.22 | 1 | 0.85 |
| ***L. peploides*** | BRT | Ensemble | Lasso |  | ***L. peploides*** | BRT | Ensemble | Lasso |
| Ensemble | 0.38 | - | - |  | Ensemble | 1 | - | - |
| Lasso | 0.38 | 0.68 | - |  | Lasso | 1 | 1 | - |
| RF | 0.42 | 1 | 1 |  | RF | 1 | 1 | 1 |
| ***L. polyphyllus*** | BRT | Ensemble | Lasso |  | ***L. polyphyllus*** | BRT | Ensemble | Lasso |
| Ensemble | 0.535 | - | - |  | Ensemble | 0.62 | - | - |
| Lasso | 0.192 | 0.535 | - |  | Lasso | 0.3364 | 0.5781 | - |
| RF | **0.021** | **0.015** | **0.015** |  | RF | **0.0221** | **0.0179** | **0.0035** |

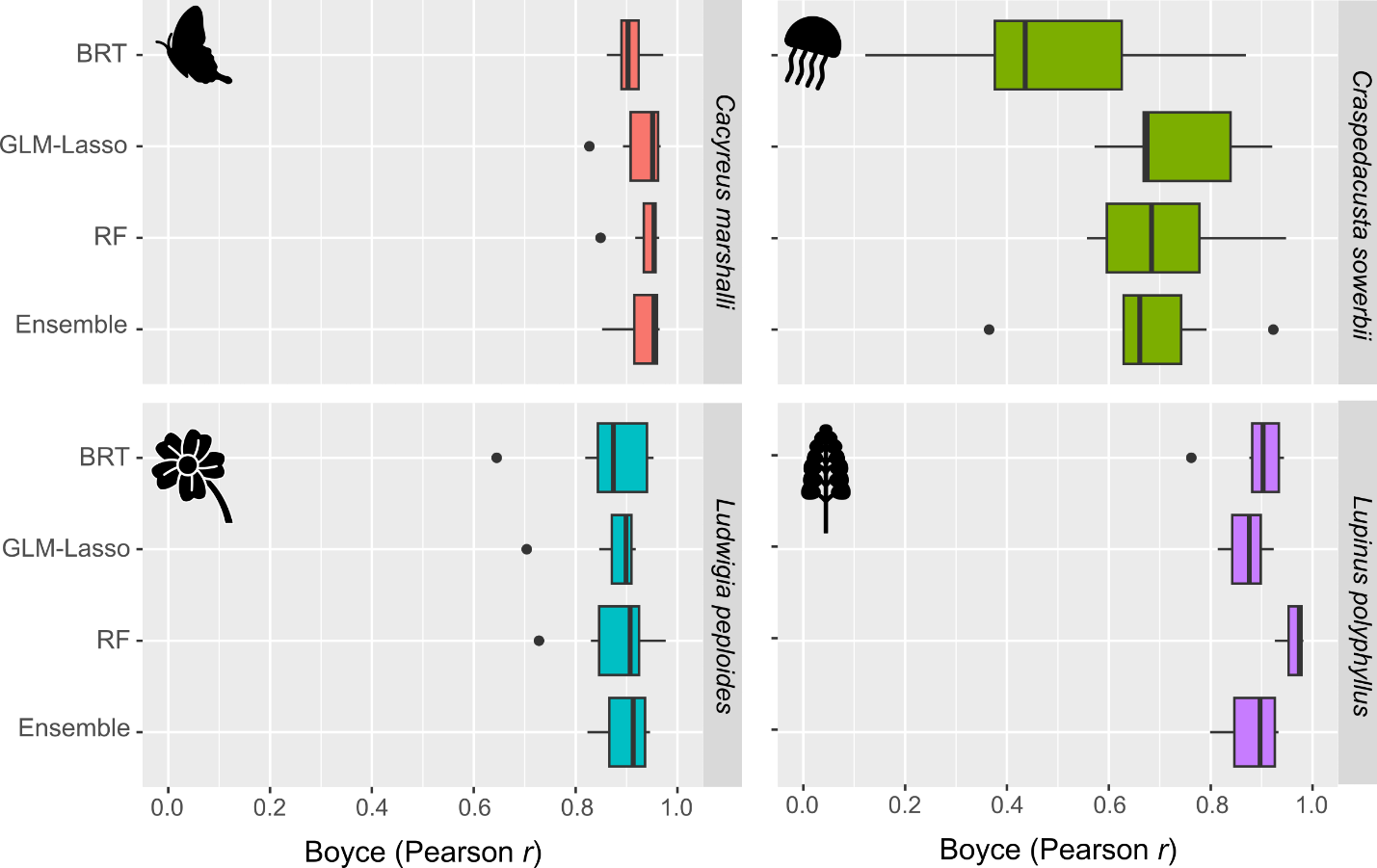

**Figure S1** – Boyce index values, corresponding to Pearson correlation values between predicted probabilities of event occurrence and the frequency of event observation records for models calibrated with data corrected for spatial and temporal bias. Four phenological stages are represented: the imago stage of the Geranium bronze butterfly (*Cacyreus marshalli*), medusae of the freshwater jellyfish (*Craspedacusta sowerbii*), and the flowering of floating primrose-willow (*Ludwigia peploides*) and the garden lupin (*Lupinus polyphyllus*). The boxplots represent the variation of correlation values assessed for 7 years (2016 to 2022), using three modelling algorithms (boosted regression trees, BRT; generalised linear models with lasso regularization, GLM-Lasso; random forest, RF) and an ensemble of previous algorithms (Ensemble), trained with observation data corrected for spatial bias.

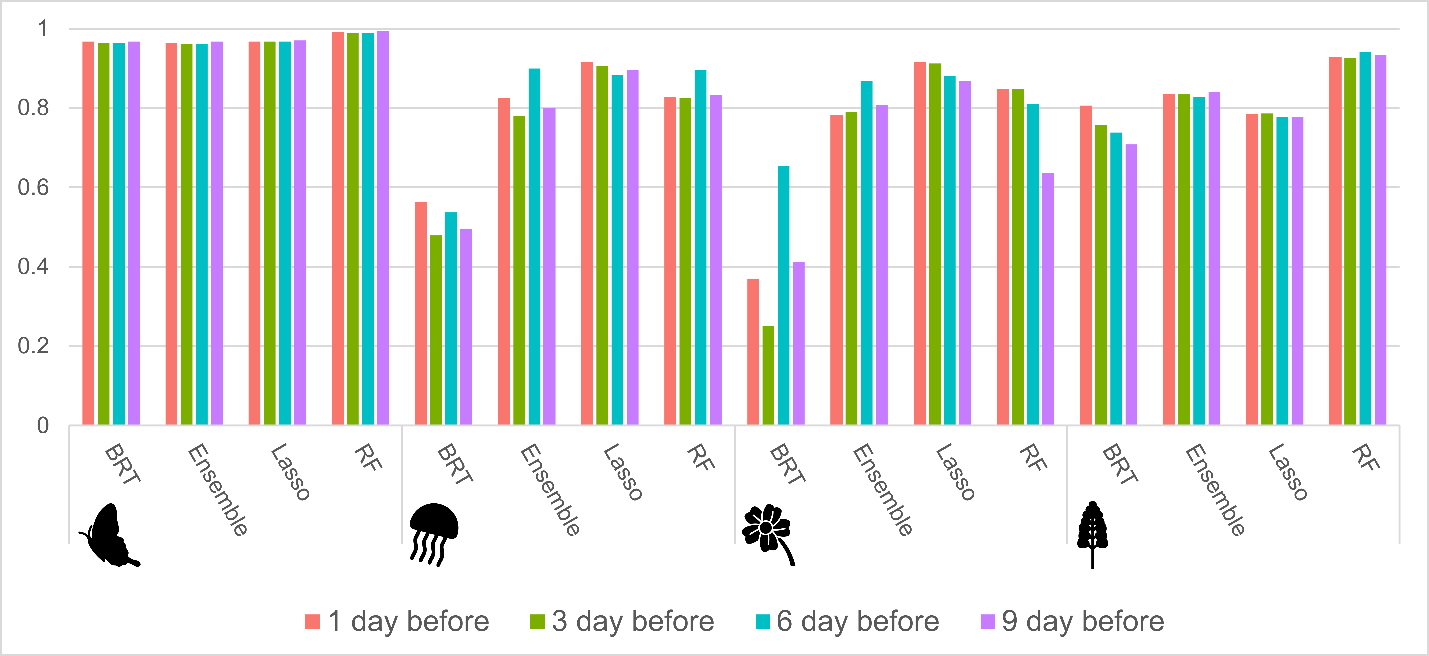

**Figure S2** – Boyce Index values for forecasts made 1, 3, 6, and 9 days in advance from models calibrated with observation data corrected for spatial and temporal bias. Represented values correspond to the Pearson correlation coefficient between predicted probabilities of event occurrence from July 2023 to March 2024 and the frequency of event observations recorded during the same period. The values are reported for three modelling algorithms—boosted regression trees (BRT), generalized linear models with lasso regularization (Lasso), and random forest (RF)—as well as an ensemble of these algorithms (Ensemble).

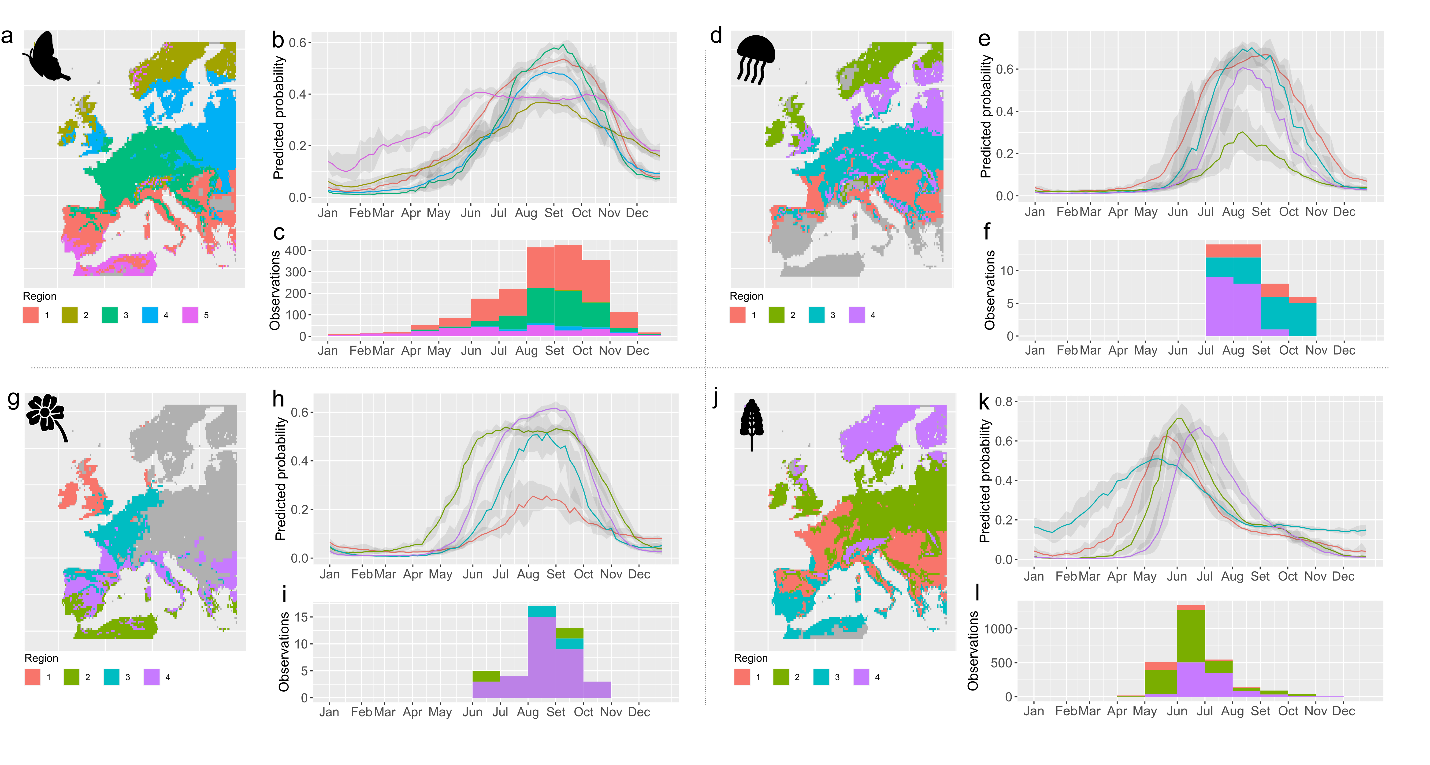

**Figure S3** – Regional patterns of predicted and observed timings of the phenological stages. Predictions are from models using observation data corrected for spatial and temporal bias. Four phenological stages are represented: the emergence of the imago stage in the Geranium bronze butterfly (*Cacyreus marshalli*; panels a-c), the occurrence of medusae in the freshwater jellyfish (*Craspedacusta sowerbii*; panels d-f), and the flowering phases of the floating primrose-willow (*Ludwigia peploides*; panels g-i) and the garden lupin (*Lupinus polyphyllus*; panels j-l). The maps display areas having similar phenological dynamics ('phenoregions'), based on daily projections at 5 days intervals from 2016 to 2022. Areas in grey represent environmental conditions far from those represented in the training data and were not considered in the analysis to minimise risks of model extrapolation. Time series depict the inter-annual mean probabilities of occurrence of each event, along with their standard deviations (grey shading), for each region throughout the year. Histograms show the monthly frequency of observed occurrences of each phenomenon within each specified region. Predictions and phenoregion delineations were made also for areas where the species have not yet been recorded, leading to observation records being absent from the histograms for certain regions.
